# Fast and accurate reference-guided scaffolding of draft genomes

**DOI:** 10.1101/519637

**Authors:** Michael Alonge, Sebastian Soyk, Srividya Ramakrishnan, Xingang Wang, Sara Goodwin, Fritz J. Sedlazeck, Zachary B Lippman, Michael C. Schatz

## Abstract

**Background:** As the number of new genome assemblies continues to grow, there is increasing demand for methods to coalesce contigs from draft assemblies into pseudomolecules. Most current methods use genetic maps, optical maps, chromatin conformation (Hi-C), or other long-range linking data, however these data are expensive and analysis methods often fail to accurately order and orient a high percentage of assembly contigs. Other approaches utilize alignments to a reference genome for ordering and orienting, however these tools rely on slow aligners and are not robust to repetitive contigs.

**Results:** We present RaGOO, an open-source reference-guided contig ordering and orienting tool that leverages the speed and sensitivity of Minimap2 to accurately achieve chromosome-scale assemblies in just minutes. With the pseudomolecules constructed, RaGOO identifies structural variants, including those spanning sequencing gaps that are not reported by alternative methods. We show that RaGOO accurately orders and orients contigs into nearly complete chromosomes based on *de novo* assemblies of Oxford Nanopore long-read sequencing from three wild and domesticated tomato genotypes, including the widely used M82 reference cultivar. We then demonstrate the scalability and utility of RaGOO with a pan-genome analysis of 103 *Arabidopsis thaliana* accessions by examining the structural variants detected in the newly assembled pseudomolecules. RaGOO is available open-source with an MIT license at https://github.com/malonge/RaGOO.

**Conclusions:** We demonstrate that with a highly contiguous assembly and a structurally accurate reference genome, reference-guided scaffolding with RaGOO outperforms error-prone reference-free methods and enable rapid pan-genome analysis.

## Background

Long read single molecule sequencing technologies commercialized by Oxford Nanopore Technologies and Pacific Biosciences have facilitated a resurgence of high-quality *de novo* eukaryotic genome assemblies [1]. Assemblies using these technologies in a variety of plant and animal species have consistently reported contig N50s over 1 Mbp, while also reconstructing higher percentages of target genomes, including repetitive sequences [2, 3]. Current long read sequencers are now able to produce over one terabase of long reads per week, presenting the opportunity for detailed pan-genome analysis of unprecedented scale and including structural variations that are notoriously difficult to detect using short read sequencing. However, lagging behind the current speed and cost of generating long read sequencing data are genome assemblers, which are still unable to resolve complex repeats and related structural variants that are widespread in eukaryotic genomes. Thus, there is a need for simplified and faster approaches to scaffold fragmented genome assemblies into chromosome-scale pseudomolecules.

Two common approaches have been used to achieve chromosome-scale assemblies, namely, reference-free (*de novo*) and reference-guided approaches. One popular reference-free scaffolding approach is to anchor genome assembly contigs to some variety of genomic map [4], such as an optical, physical or linkage map [5]. This process involves aligning the genomic map to a sequence assembly and scaffolding contigs according to the chromosomal structure indicated in the map. However, contigs not implicated in any alignments will fail to be scaffolded, which can result in incomplete scaffolding. Furthermore, acquiring a genomic map can be expensive, time consuming, or otherwise intractable depending on the species and the type of map.

Another reference-free method for pseudomolecule construction involves the use of long-range genomic information to scaffold assembled contigs. This includes a large class of technologies such as mate-pair sequencing, Bacterial Artificial Chromosomes (BACs), Linked Reads and chromatin conformation [6-8]. In particular, Hi-C has recently been shown to a be a practical and effective resource for chromosome-scale scaffolding [9-11]. Paired-end Hi-C sequencing reads are aligned to the assembly and mates which align to different contigs (Hi-C links) are recorded. According to the relative density of such Hi-C links between pairs of contigs, contigs can be ordered and oriented into larger scaffolds, potentially forming chromosome-length pseudomolecules. Also, because misassemblies may be observed by visualizing Hi-C alignments, Hi-C can be used for validation and manual correction of misassemblies[12]. Though Hi-C has been widely adopted, there remain challenges that can impede the ability to form accurate chromosome-scale pseudomolecules with Hi-C alone. Principally, Hi-C data are noisy, and Hi-C based scaffolders are prone to producing structurally inaccurate scaffolds [13]. Also, because this process relies on alignment of short Hi-C sequencing reads to the draft assembly, small and repetitive contigs with little or conflicting Hi-C link information often fail to be accurately scaffolded. Finally, the analysis requires deep sequencing coverage and therefore can be expensive and compute intensive.

Aside from reference-free approaches to scaffolding, tools such as Chromosomer and MUMmer’s ‘show-tiling’ have been developed for reference-guided pseudomolecule construction [14-17]. Such tools utilize alignments between a genome assembly and a closely related reference assembly for scaffolding. Similarly, other tools use multiple, potentially diverse references for scaffolding [18, 19]. Though reference-guided scaffolding may introduce erroneous reference-bias, it is often substantially faster and less expensive than acquiring the resources for the reference-free methods outlined above. However, current tools for reference-guided scaffolding of eukaryotic genomes have notable shortcomings. Firstly, these tools depend on slower DNA aligners such as BLAST, Nucmer and Cactus, and accordingly require long compute times of several hours to several days for mammalian-sized genomes [20-22]. These aligners are also not robust to repetitive and/or gapped alignments resulting in a significant portion of contigs being unlocalized in pseudomolecules. Finally, many of these methods do not internally offer the ability to correct large-scale misassemblies frequently present in draft assemblies of eukaryotic genomes nor report any metrics on conflicts due to true biological differences in the genomes.

Here, we introduce RaGOO, an open-source method which utilizes Minimap2 [23] alignments to a closely related reference genome to quickly cluster, order and orient genome assembly contigs into pseudomolecules. RaGOO also provides the option to correct apparent chimeric contigs prior to pseudomolecule construction. Finally, structural variants (SVs), including those spanning gap sequence, are identified using an optimized and integrated version of Assemblytics [24], thus enabling rapid pan-genome SV analysis of many genomes at once. This is especially important for detecting large insertions and other complex structural variations that are difficult to detect using read mapping approaches.

We first demonstrate the speed and accuracy of RaGOO scaffolding with simulated data of increasing complexity and show that it outperforms two popular alternative methods. We next show the utility of RaGOO by creating high quality chromosome-scale reference genomes for three distinct wild and domesticated genotypes of the model crop tomato using a combination of short and long read sequencing. Finally, we demonstrate the scalability of RaGOO by ordering and orienting 103 draft *A. thaliana* genomes and comparing structural variants across the pan-genome. This uncovers a large number of defense response genes that are highly variable.

## Results

### Reference-guided Contig Ordering and Orientation with RaGOO

RaGOO is an open-source tool, implemented as a python command-line utility, to order and orient genome assembly contigs according to Minimap2 alignments to a single reference genome (**Figure 1**). RaGOO’s primary goal is borrow the large-scale structure of a reference genome to organize assembly contigs, analogous to how a genetic map is used. Therefore, under default settings, RaGOO does not alter or mutate any input assembly sequence, but rather arranges them and places gaps for padding between contigs. Additionally, users have the option to break input contigs at points of misassembly indicated by non-sequential alignments to the reference genome. However, these breaks can only further fragment the assembly and do not add or remove any sequence content. RaGOO can optionally avoid breaking chimeric intervals at loci within genomic coordinates specified by a gff3 file, such as to avoid disrupting gene models identified in the de novo assembly. Additionally, RaGOO computes confidence scores associated with the clustering, ordering and orienting of contigs. These scores can be used to assess how scaffold localization is supported by the underlying alignments and can be helpful in determining the success of chimeric contig correction. This can also be used to help identify true biological differences between the reference genome and the newly assembled sample.

**Figure 1.**
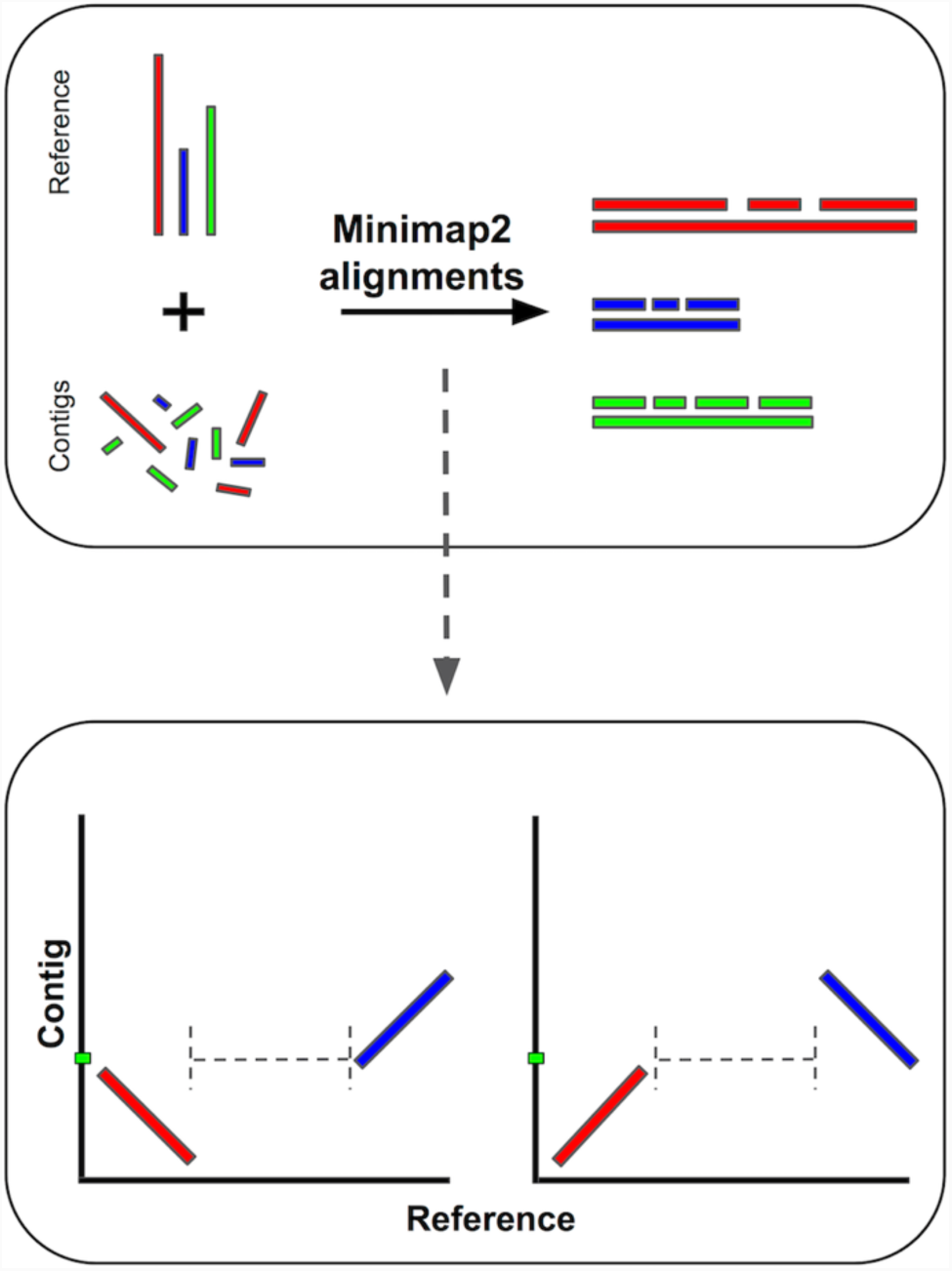
The RaGOO Pipeline. **(top)** Contigs are aligned to the reference genome with Minimap2, and are ordered and oriented according to those alignments. Bottom: Optionally, chimeric contigs, contigs with large distances (dashed lines) between consecutive alignments (blue and red lines), can be broken prior to ordering and orientation. Green bars represent contig breakpoints.

After constructing pseudomolecules, with or without chimeric contig correction, RaGOO re-aligns the assembly to the reference and calls structural variants with an integrated version of Assemblytics. We have optimized this approach by replacing the relatively slow single-threaded nucmer alignment phase with the much faster Minimap2 aligners along with the necessary converters between the output formats. Noting that such alignments may traverse gaps in either the reference or the assembly, we report the percent overlap between each SV and gaps, allowing users to utilize such variants at their discretion. Importantly, the speed of Minimap2 alignments, and therefore RaGOO, facilitates a genome scaffolding and SV analysis at scales previously not feasible with comparable tools. For example, RaGOO scaffolds an *Arabidopsis thaliana* draft assembly (7,472 total contigs) in 73 seconds as a single process on a MacBook Pro laptop.

### Simulated Reference-Guided Scaffolding

To assess the efficacy of RaGOO, we used it to scaffold simulated draft eukaryotic genome assemblies of increasing difficulty. To simulate these assemblies, we partitioned the current tomato (*Solanum lycopersicum*) reference genome (Heinz version SL3.0) into variable length scaffolds [25]. To achieve a realistic distribution of sequence lengths, we sampled the observed contig lengths from a *de novo* assembly produced with Oxford Nanopore long reads of the M82 *S. lycopersicum* cultivar, which is described later in this paper **(Methods**). Given that many of these resulting scaffolds contained gap sequence from the reference genome, we also established an assembly comprised of contigs free of sequencing gaps (“N” characters). For this, we split the simulated scaffolds at any stretch of 20 or more “N” characters, excluding the gap sequence. We also excluded any resulting contigs shorter than 10 kbp in length. We refer to these scaffolds and contigs as the “easy” set of simulated data, as they are a partitioning of the reference with no variation. To simulate a “hard” dataset that contained variation, we used SURVIVOR [26] to simulate 10,000 insertion and deletion SVs, ranging in size from 20 bp to 10 kbp in size, and SNPs at a rate of 1% into the simulated scaffolds. Contigs were then derived from these scaffolds just as with the contigs. Assembly stats for these 4 simulated assemblies are in **Supplementary Table 1**.

Utilizing the same SL3.0 reference assembly, we then used MUMmer’s ‘show-tiling’ utility, as well as Chromosomer and RaGOO to arrange these simulated assemblies into 12 pseudomolecules. To assess scaffolding success, we measured clustering, ordering and orienting accuracy. Clustering and orienting accuracy is the percentage of localized contigs that were assigned the correct chromosome group and orientation respectively. To assess ordering accuracy, the edit distance between the true and predicted contig order was calculated for each pseudomolecule normalized by the true number of contigs in the pseudomolecule. Additionally, for a local measurement of ordering accuracy, the fraction of correct adjacent contig pairs was computed for each pseudomolecule. Finally, to measure scaffolding completeness, we noted the percentage of contigs and total sequence localized into pseudomolecules.

RaGOO performed well on all datasets, achieving high clustering, ordering and orienting accuracy on both the “easy” and “hard” datasets, while localizing the vast majority (∼99.9998% for hard scaffolds) of sequence in only a few minutes (∼2 minutes for the “hard” scaffolds) (**Figure 2, Supplementary Table 2**). In all simulations, Chromosomer accurately reconstructed most of the genome, though the presence of gaps in scaffolds and variation in the “hard” assembly degraded performance to a localization score of 86.65% in the “hard” scaffolds. Show-tiling suffered tremendously from the presence of gaps in scaffolds and accordingly achieved poor localization scores on scaffolds of both the “easy” (8.43%) and “hard” (0.01%) sets. Both Chromosomer and show-tiling took substantially longer to run than RaGOO in all cases, completing in several hours rather than minutes.

**Figure 2.**
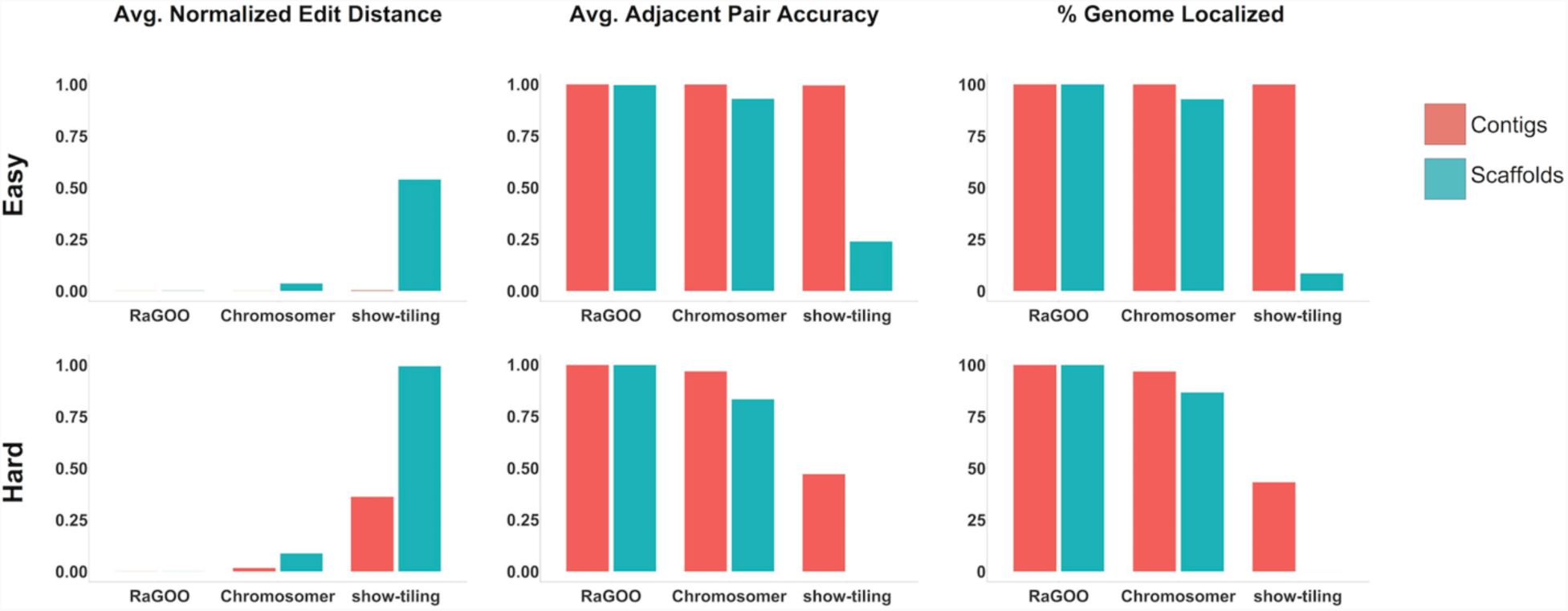
Scaffolding Simulated Assemblies. Ordering and localization results for “easy” and “hard” simulated tomato genome assemblies. Normalized edit distance and adjacent pair accuracy measure the success of contig ordering and are averaged across the 12 simulated chromosomes. The percentage of the genome localized measures how much of the simulated assemblies were clustered, ordered and oriented into pseudomolecules.

### Pan-SV Analysis of 3 Chromosome-scale Tomato Genome Assemblies

For more than a decade, the reference genome for tomato (var. ‘Heinz 1706’) has been an invaluable resource in both basic and applied research, but extensive sequence gaps (81.7 Mbp, 9.87%), unlocalized sequence (∼17.8 Mbp, 2.39%), and limited information on natural genetic variation in the wider germplasm pool impeded its full utilization [25]. To compensate, more than 700 additional accessions have since been sequenced by Illumina short-read technology [27, 28]. However, due to the short sequence reads these studies were limited to evaluating, with reasonable accuracy, (depending on variable sequencing quality and coverage) single nucleotide polymorphisms (SNPs) and small insertions and deletions (indels). In contrast, larger structural variations (SVs) that have important and often underestimated functional consequences for genome evolution and phenotypic diversity were largely ignored in this major model crop plant. Critically, without long reads, the complete catalog of structural variations in the species, a pan-SV analysis, is largely incomplete.

To address this knowledge gap and begin constructing a high-quality tomato pan-SV analysis, we used long-read Oxford Nanopore Technology (ONT) to sequence three distinct genotypes that provide anchor points for wild and domesticated tomato germplasm: 1) the species *S. pimpinellifolium* is the ancestor of tomato, and the Ecuadorian *S. pimpinellifolium* accession BGV006775 (BGV) represents the group of progenitors that are most closely related to early domesticated types; 2) The processing cultivar *S. lycopersicum* M82 is the most widely used accession in research due to its rich genetic resources, and 3) the elite breeding line *S. lycopersicum* Fla.8924 (FLA) is a large-fruited ‘fresh-market’ type that was developed for open field production in Florida [29, 30]. Together, these three accessions provide a foundation for constructing a pan-SV analysis that will enable identification and classification of thousands of predicted SVs.

#### Reference-guided and reference-free M82 scaffolding

In order to evaluate the effectiveness of RaGOO with genuine sequencing data, we first used it along with other reference-guided and reference-free tools to scaffold a highly contiguous assembly of the M82 *S. lycopersicum* cultivar. We sequenced the genome with an Oxford Nanopore MinION sequencer to 58.8x fold coverage with an N50 read length of 13.4kbp (max: 1,256,650bp). The genome was assembled with Canu [31] and was comprised of 1,709 contigs with a contig N50 of 1,458,445 bp. To compare RaGOO to other reference-guided tools, the assembly was scaffolded with RaGOO (with chimeric contig correction), MUMmer’s ‘show-tiling’ utility, and Chromosomer. Here, a “localized” contig is one that is placed in a pseudomolecule group and is assigned an order and orientation. In all cases, the Heinz SL3.0 genome was used as the reference. RaGOO localized the highest portion of sequence, placing 99.01% of sequence into chromosomes compared to 85.6% and 3.17% for Chromosomer and show-tiling respectively (**Supplementary Table 3**). The resulting RaGOO assembly contained 12 chromosome-length pseudomolecules with only 0.99% of sequence in the ambiguous chromosome 0 (**Figure 3**). Additionally, the scaffolding completed in only ∼7 minutes for RaGOO, compared to ∼285 minutes for show-tiling and ∼1,466 minutes for Chromosomer.

**Figure 3.**
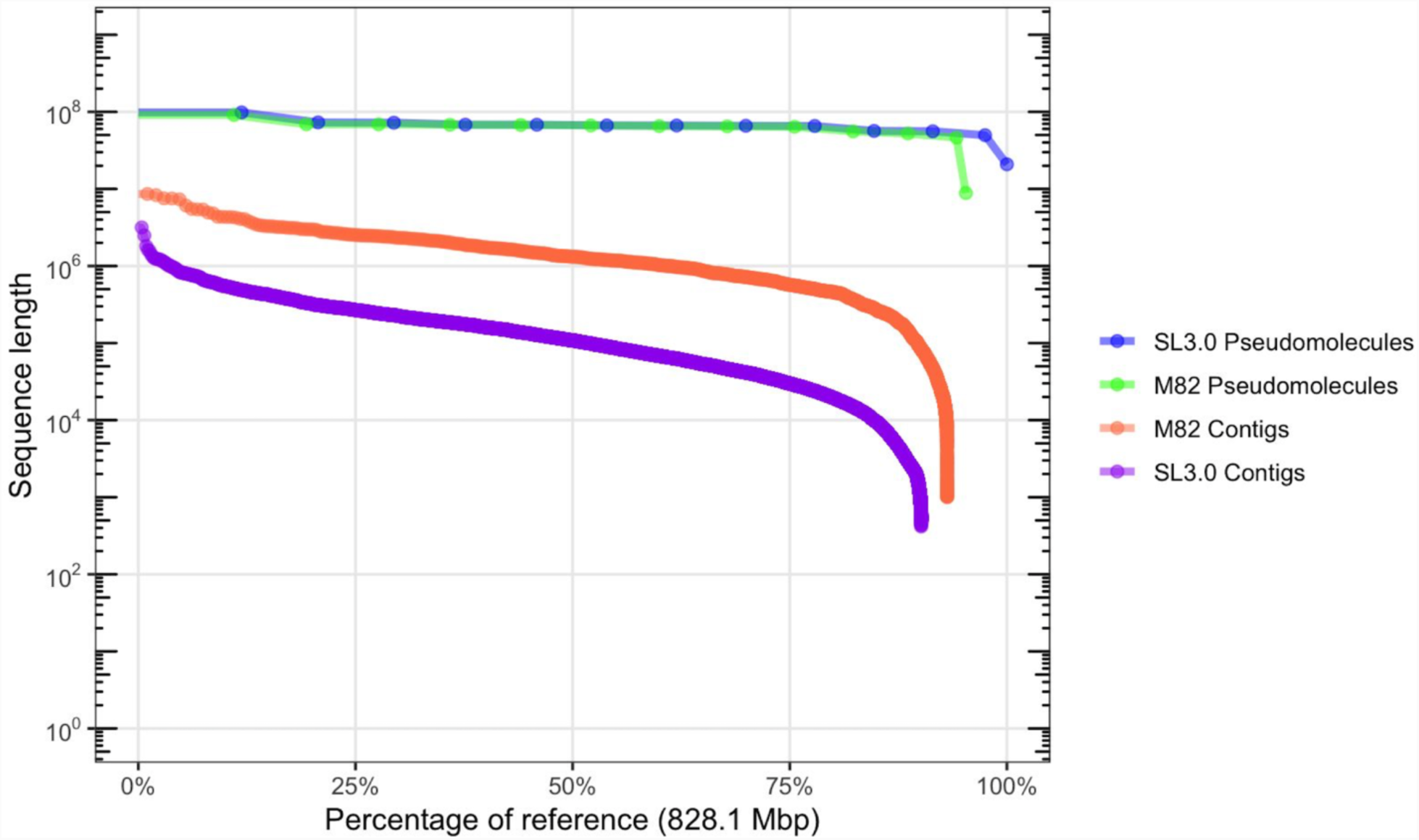
M82 Assembly Contiguity. “Nchart” of the M82 and Heinz contigs and pseudomolecules. M82 pseudomolecules were established by ordering and orienting M82 contigs with RaGOO. Heinz contigs were derived from the SL3.0 pseudomolecules by splitting sequences at stretches of 20 or more contiguous “N” characters.

To compare RaGOO scaffolding to a widely used reference-free approach, we generated Hi-C chromatin conformation data and used SALSA2 [13] to build scaffolds from the M82 contigs. Though SALSA2 does not necessarily build pseudomolecules, it strives to establish chromosome and chromosome-arm length scaffolds as the data allows. SALSA2 utilized Hi-C alignments to the M82 draft assembly along with the M82 Canu assembly graph. Though the scaffolds were highly contiguous compared to the input assembly (scaffold N50 of 18,282,950 bp), they fall far short of complete chromosome-scale.

We further compared the structural accuracy of the RaGOO pseudomolecules to that of the SALSA2 scaffolds by comparing the 12 pseudomolecules of the former and the 12 longest scaffolds of the latter to the Heinz SL3.0 reference. The dotplots from these alignments are displayed in **Figure 4 (left)**. This shows nearly complete and highly co-linear coverage of the RaGOO pseudomolecules, while highly fragmented and rearranged placements of the SALSA2 scaffolds. Additionally, realigning the same Hi-C data to these pseudomolecules/scaffolds provides a reference-free assessment of the large-scale structural accuracy of these sequences. Through this analysis, we found that the SALSA2 scaffolds contained many misassemblies, especially false inversions, while the RaGOO pseudomolecules contained very few structural errors (**Figure 4 right**). These Hi-C alignments suggest that most inversions and other large structural differences between the SALSA2 scaffolds and the Heinz reference assembly are likely not biological, but rather, are scaffolding errors. They also demonstrate that erroneous reference-bias in the RaGOO pseudomolecules, though present, was rare.

**Figure 4.**
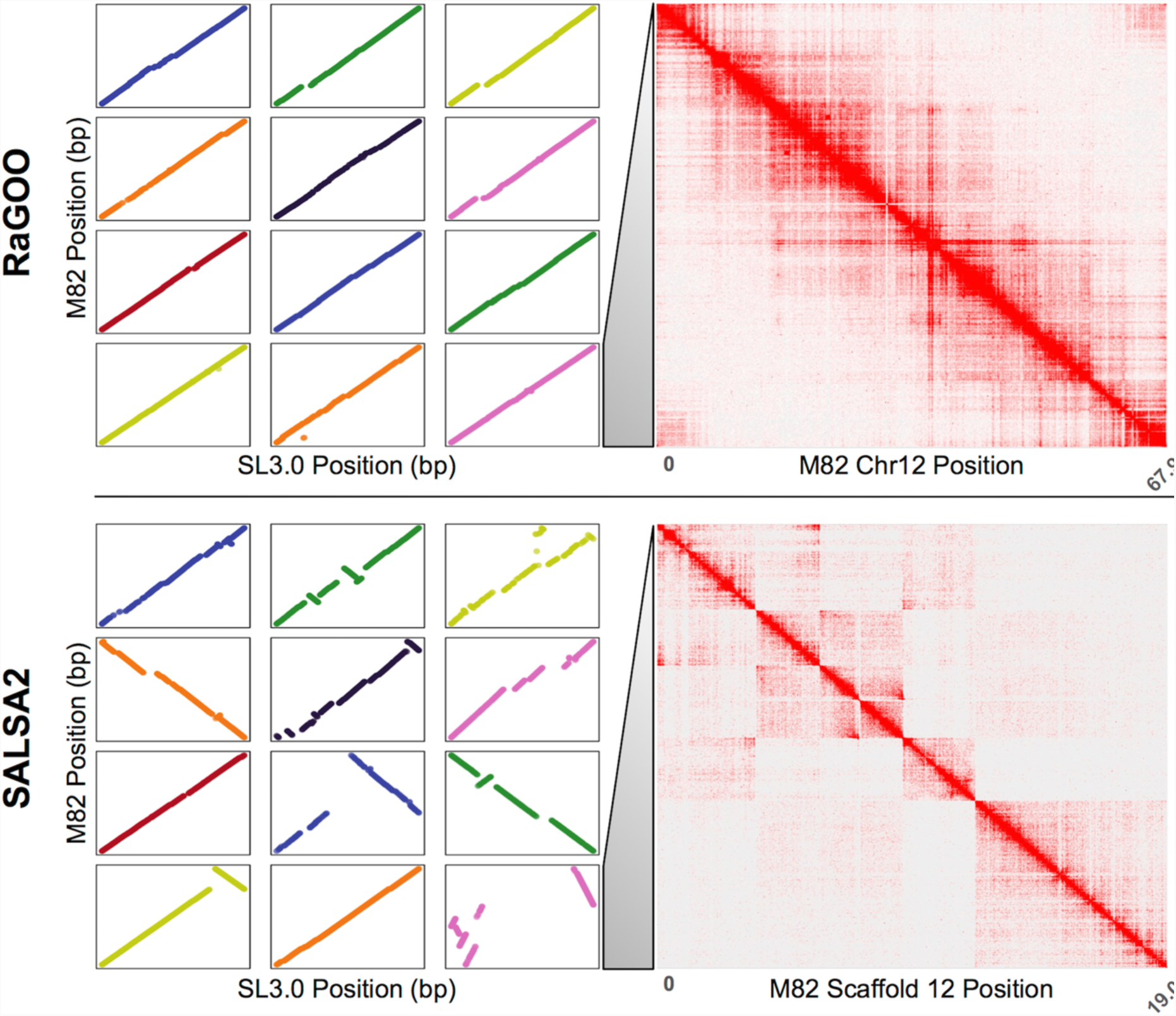
Reference-Free vs. Reference-Guided Scaffolding of M82. Both the top and bottom panels depict a dotplot (left) and Hi-C heatmap (right). The dotplots are generated from alignments to the Heinz reference assembly. On the top panel is the reference-guided RaGOO assembly dotplot, with chromosomes 1 through 12 depicted from top left to bottom right, and the Hi-C heatmap for chromosome 12. On the bottom is the *de novo* SALSA scaffolds dotplot, with the 12 largest scaffolds depicted in descending order of length from top left to bottom right, and the Hi-C heatmap for the 12th largest scaffold.

#### M82 Chromosome Hi-C validation and finishing and annotation

In an effort to establish a new structurally accurate tomato reference genome, we used the Hi-C data and Juicebox Assembly Tools to correct apparent misassemblies starting with the RaGOO M82 pseudomolecules as they provided the best completeness and contiguity with relatively few misassemblies. A total of 3 corrections were made: an inversion error correction on chromosome 3 and an ordering error correction on chromosome 7 and 11. Any “debris” contigs resulting from these alterations were placed in chromosome 0. With these few misassemblies corrected, the pseudomolecules were gap filled with PBJelly and polished with Pilon [32, 33] (**Methods**, **Supplementary Table 4**). The final polished assembly had an average identity of 99.56% when comparing to the Heinz SL3.0 reference and contained a complete single copy of 94.1% of BUSCO genes [34]. We note that M82 is biologically distinct from Heinz so we do not expect 100% identity and estimate the overall identity at approximately 99.8% to 99.9%. Additionally, M82 consensus accuracy is reflected in ITAG 3.2 cDNA GMAP alignments, 96.8% of which align with at least 95% coverage and identity (**Supplementary Figure 1**) [35].

Gene finding and annotation was performed on the finished M82 assembly with the MAKER pipeline [36] (**Methods**, **Supplementary Figure 2**, **Supplementary Table 5**). There are 35,957 genes annotated in the M82 assembly, of which 27,624 are protein coding. When comparing M82 and Heinz 1706 ITAG3.2 gene models using gffcompare (https://github.com/gpertea/gffcompare), we found 24,652 gene models with completely matching intron chains. The final M82 assembly contained a total of ∼46 Mbp novel non-gapped sequence missing from the SL3.0 reference genome. Furthermore, the M82 assembly contained only ∼8.9 Mbp of unlocalized sequence in chromosome 0 compared to ∼ 17.8 Mbp in the Heinz SL3.0 reference.

#### Pan-SV analysis of 3 tomato accessions

In addition to the M82 cultivar, we also assembled genomes for the BVG and FLA tomato accessions *de novo* with Oxford Nanopore sequencing reads and the Canu assembler. We sequenced the BGV accession to 33.5x fold coverage with a read N50 length of 27,350bp (max: 192,728bp), and the FLA accession to 41.6x fold coverage with a read N50 length of 24,225bp (max: 144,350bp). The FLA assembly contained a total of 750,743,510 bp and had an N50 of 795,751 bp, while the BGV assembly contained a total of 769,694,915 bp and had an N50 of 4,105,177 bp. As with the M82 assembly, RaGOO was then used to establish pseudomolecules and call structural variants for these assemblies. The final FLA and BGV pseudomolecules contained 745,663,382bp and 765,377,903bp (99.3% and 99.4%) of total ungapped sequence localized to chromosomes respectively. Finally, the assemblies underwent gap filling, polishing and gene finding using the same methods as M82 (**Supplementary Table 5**). A summary of the final assembly statistics for all three accessions are presented in **Table 1**. The polished assemblies had 99.4% (FLA) and 98.9% (BGV) average identity compared to the Heinz SL3.0 reference as measured by MUMmer’s ‘dnadiff’. These assemblies also demonstrated genome completeness with BGV containing a single copy of 94.8% and FLA containing a single copy of 94.9% of BUSCO genes.

**Table 1.** Summary statistics of the reference tomato genome as well as the 3 novel accessions. Chromosome span indicates total span of all of the chromosomes, including gaps. Chromosome N50 is the length such that half of the total span is covered in chromosome sequences this length or longer. Chr0 bases reports the number of bases assigned to the unresolved Chromosome 0. Contig Span is the total length of non-gap (N) characters. Contig N50 is the length such that half of the contig span is covered by contigs this length or longer. Number SVs reports the number of SVs reported by RaGOO using the integrated version of Assemblytics.

| Accession | Chromosome Span (bp) | Chromosome N50 (bp) | Chr0 bases (bp) | Number Contigs | Contig Span (bp) | Contig N50 (bp) | Number SVs |
| --- | --- | --- | --- | --- | --- | --- | --- |
| **Heinz** | 828,076,956 | 66,723,567 | 20,852,292 | 22,705 | 746,357,581 | 133,084 | NA |
| **M82** | 792,934,937 | 67,021,692 | 8,891,603 | 2,910 | 771,143,786 | 1,458,445 | 36,191 |
| **BGV** | 794,568,563 | 67,174,401 | 4,643,553 | 638 | 769,694,915 | 4,105,177 | 45,927 |
| **FLA** | 796,004,315 | 67,650,907 | 5,490,904 | 2,577 | 750,743,510 | 795,751 | 45,478 |

Together with the M82 genome, we present 3 chromosome-scale assemblies with substantially more sequence content and fewer gaps than the Heinz SL3.0 reference genome. Given structural variants output by RaGOO, we next used SURVIVOR to determine which variants were shared amongst these three accessions (**Figure 5**). As expected, the most divergent accession, BGV, demonstrated the most structural variant diversity with a total of 45,927 SVs compared to 45,478 and 36,191 SVs in FLA and M82 respectively. The union of these sets of variants yielded 98,988 total structural variants, which overlapped with 19,790 out of 35,768 total ITAG 3.2 genes (with 2 kbp flanking upstream and downstream each gene included). A complete list of gene/variant intersections is available in **Supplementary Table 6**. The most variable gene (the gene with the most intersecting SVs), Solyc03g095810.3, is annotated as a member of the GDSL/SGNH-like Acyl-Esterase family, while the second most variable gene, Solyc03g036460.2, is annotated as a member of the E3 ubiquitin-protein ligase. These three chromosome-scale assemblies, along with their associated sets of SVs, establish valuable genomic resources for the *Solanaceae* scientific community.

**Figure 5.**
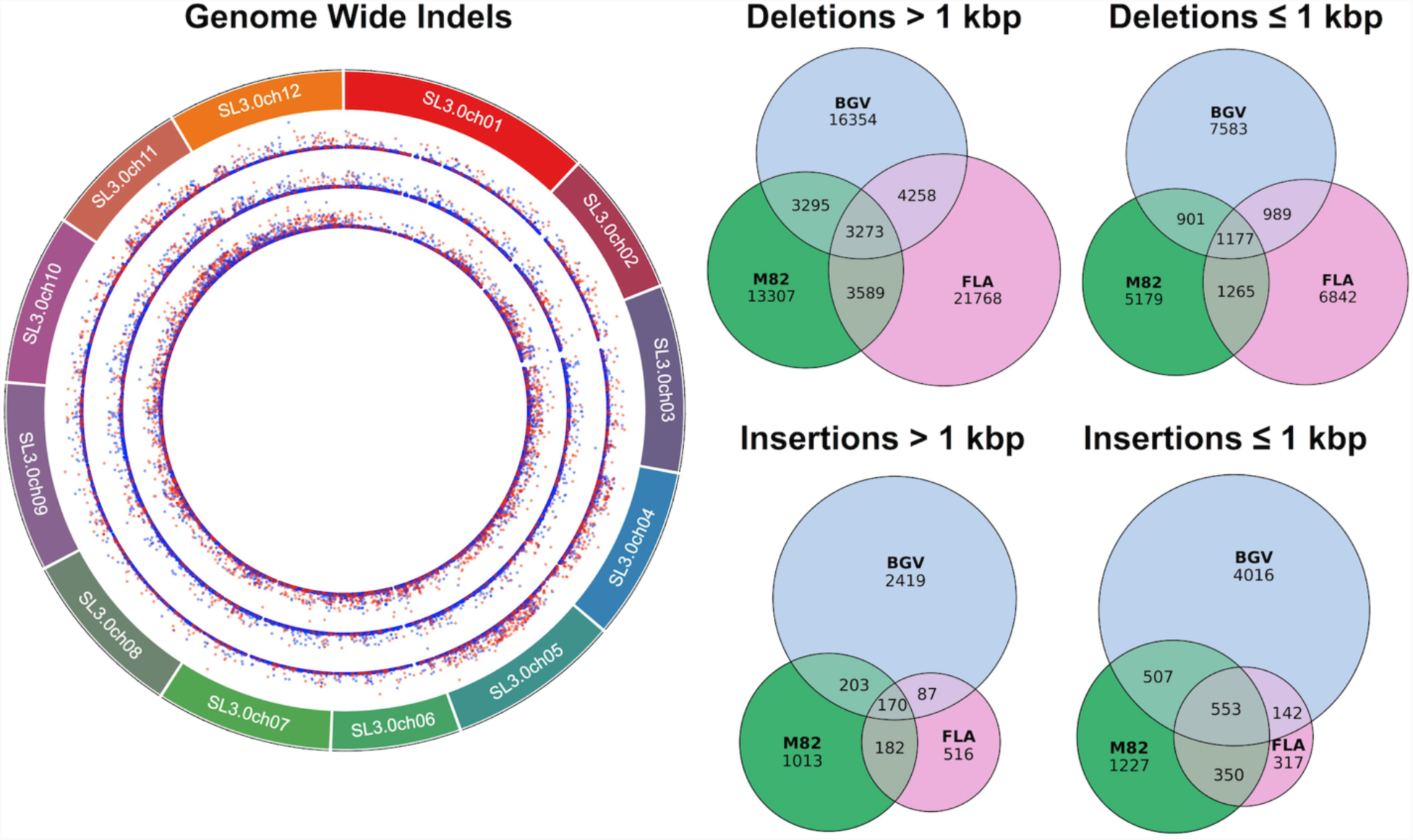
The Tomato Pan-Genome. **(left)** Circos plot (http://omgenomics.com/circa/) depicting size and type of structural variant. From outer ring to inner ring: M82, FLA and BGV. Point height (y-axis) is scaled by the size of the variant, with red indicating insertions and blue indicating deletions. **(right)** Euler diagrams (https://github.com/jolars/eulerr) depicting insertions and deletions shared amongst the three accessions.

### Pan-SV analysis of 103 *Arabidopsis thaliana* genomes

Given the speed of RaGOO, we sought to test its performance power by performing a panSV-genome analysis on a large population of diverse individuals. To acquire such population-scale data, we examined sequencing data from the 1001 genomes project database, which includes raw short-read sequencing data and small variant calls for 1,135 *Arabidopsis thaliana* accessions [37]. We mined the 1001 genomes project database for sequencing data amenable to genome assembly with sufficiently deep coverage of paired-end reads (**Methods**). This identified 103 short-read datasets representing a wide range of accessions sampled across 4 continents (**Figure 6.A**). We then established draft *de novo* assemblies for each accession using SPAdes [38]. Finally, RaGOO utilized the TAIR 10 reference genome to create 103 chromosome-scale assemblies and associated SV calls [39]. Between 85.8% and 98.7% (mean=96.7%) of sequence was localized into chromosomes per accession, showing that the majority of assembled sequence across the pan-genome was scaffolded into pseudomolecules, even for more divergent accessions. The structural variant calls from this pan-genome provide a database of *A. thaliana* genetic variation previously unreported in the initial 1001 genomes project analysis [40].

**Figure 6.**
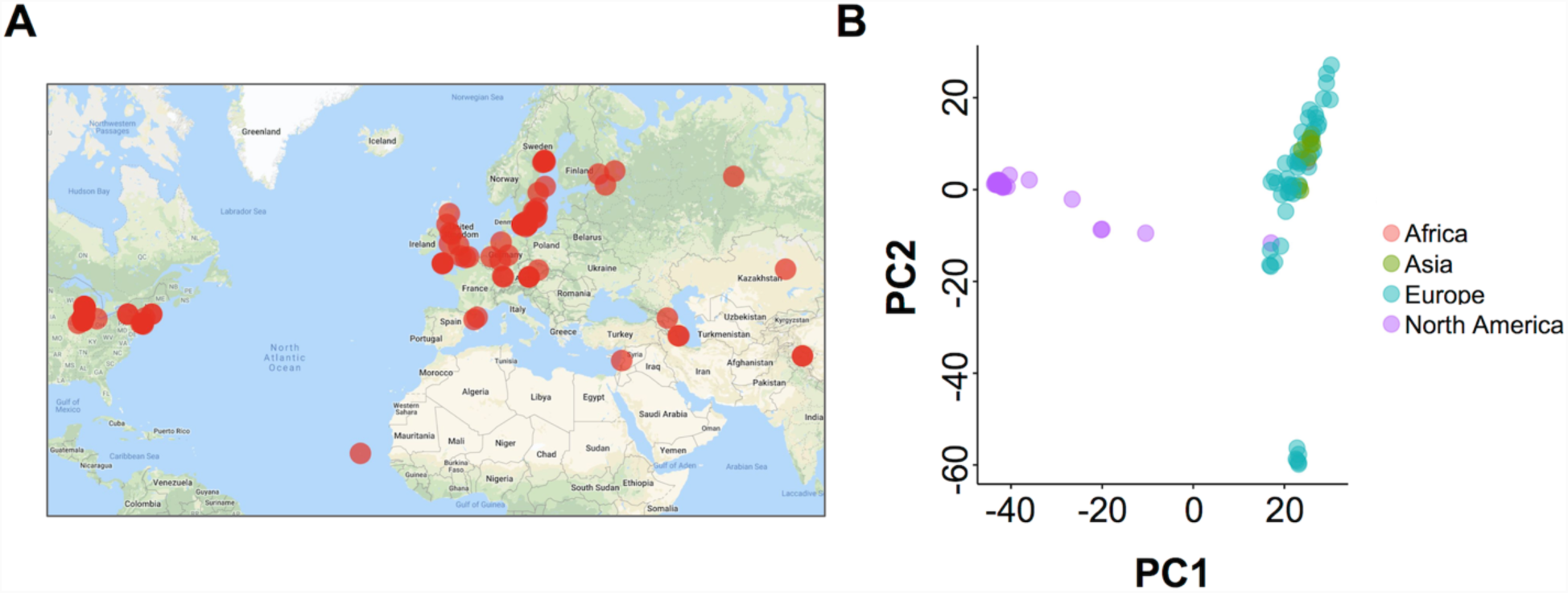
The *Arabidopsis* Pan-Genome. **(A)** Map of the 103 *Arabidopsis* accessions that were assembled in this study. **(B)** Principal components analysis of the structural variant presence/absence matrix of the 103 Arabidopsis accessions.

SV calls were compared with SURVIVOR, yielding a total of 137,111 merged variants across the pan-genome. From this merged set of variants, we constructed a presence/absence matrix representing which variants were present in which accessions. Principal Components Analysis of this matrix revealed a clustering of accessions according to their geographic location (**Figure 6.B**). Upon further analysis of global trends in the data, we found that SVs were concentrated in pericentromeric regions, consistent with previous findings (**Supplementary Figure 3**) [41].

We further examined those genes that intersected variants present in small and large numbers of accessions, as these represent rare variants in the population and rare variants in the reference genome, respectively. When including variants present in at least 1, 10, 50 and 100 samples, we found 26,795, 17,593, 7,859 and 332 total intersecting protein coding genes (2 kbp flanking each side) respectively. Since there are a total of 27,416 protein coding genes in the TAIR 10 database, we conclude that SVs in the pan-genome impact the genomic architecture for the majority of protein coding genes, though fewer genes are affected by variants present in multiple samples. The full catalog of the gene structural variations is presented in **Supplementary Table 7,** and the ten most frequently affected genes are presented in **Table 2.** Interestingly, most of these highly variable genes are defense response genes. Ultimately, our analysis highlights the importance of chromosome-level assembly at population scale to uncover genomic variation not discovered through traditional short-read mapping.

**Table 2.**
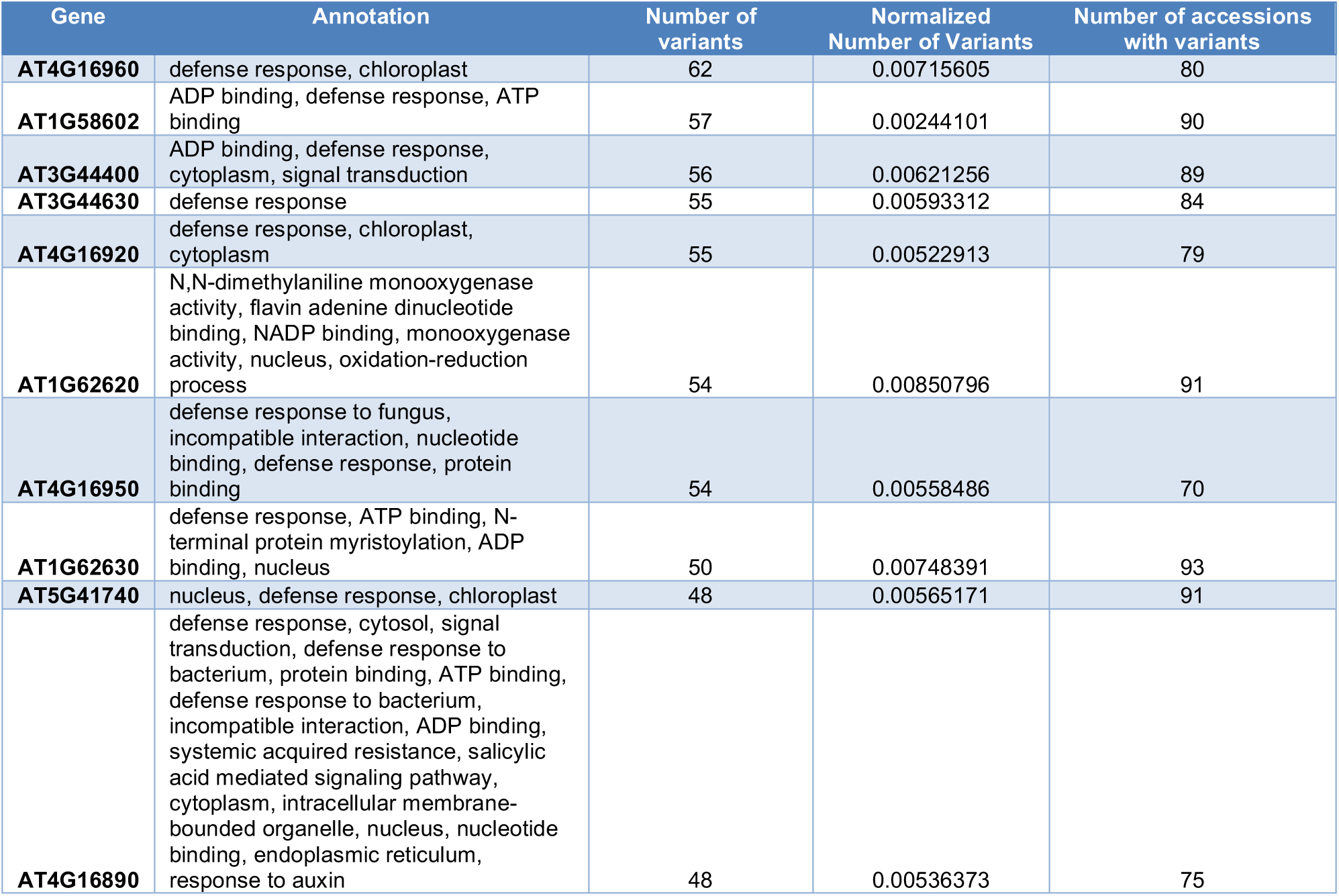
Summary of the ten most variable genes in the Arabidopsis pan-genome. “Number of variants” is the total number of variants intersecting a given gene, and “Normalized Number of Variants” is the number of intersecting variants divided by gene length.

## Discussion

We have introduced RaGOO in both a general and focused context for highly accurate genome scaffolding. As a general method, RaGOO may be valuable for chromosome-scale scaffolding in experimental designs where ordering and/or orienting of contigs leveraging an existing reference is available. Ordering and orienting with RaGOO may also facilitate analysis not possible with unlocalized contigs. This is exemplified by the additional sequence found through gap filling of the M82, BGV, and FLA assemblies or by the identification of structural variants spanning gaps between contigs in the *S. lycopersicum* and *Arabidopsis thaliana* pan-genomes. Additionally, our pan-genome analysis demonstrates that the speed of RaGOO offers new possibilities as to the scope and size of experiments that require reference-guided scaffolding. Furthermore, the integrated structural variant identification pipeline allows for a rapid survey of gene related and other variants in the population. This shows that for both tomato and *Arabidopsis* pan-genomes, the majority of protein coding genes are associated with structural variation, highlighting the importance of population-scale assembly and structural variant discovery.

In a more focused analysis, we demonstrate that RaGOO may be a valuable component of a detailed assembly pipeline to establish new high-quality eukaryotic genomic resources. Our use of RaGOO to produce three tomato assemblies highlights a valuable means of organizing contiguous draft assemblies into pseudomolecules. This is especially useful as draft assemblies become more contiguous and high-quality references become more common, even for non-model species.

Regardless of the application, there are a few key considerations to be made when using RaGOO or other reference-guided assembly/scaffolding tools. Reference-guided pseudomolecule structural accuracy is dependent on reference accuracy, draft assembly accuracy, and conservation of chromosomal structure between the reference and draft assembly genotypes. Accurate reference and draft assemblies ensure that discrepancies between the two sequences are likely biological, and contiguous draft assemblies are needed to contain the majority of biological variation within contigs. Therefore, if the large-scale chromosomal structure is also conserved between assemblies, we assert that most biological variation will not be lost during the ordering and orienting process. In the case of the M82 assembly, we showed that these aforementioned conditions were favorable, especially by showing that the Hi-C data validates the large-scale structural accuracy of the M82 pseudomolecules. Notwithstanding these positive results, additional independent information provided by Hi-C analysis can be used to correct the few remaining instances of misassembly and reference bias. From a utilitarian perspective, the reference-guided RaGOO scaffolds required less manual correction of errors than the reference-free scaffolds, and accordingly were the best option for downstream analysis.

For applications that do not have independent data such as Hi-C to validate the accuracy of RaGOO output, it is challenging to assess the extent to which errors such as reference bias are present in pseudomolecules. However, it is possible to estimate the fidelity of newly created pseudomolecules to the reference. This can be done by observing the percentage of localized contigs/sequence along with the RaGOO confidence scores. These confidence scores, described in the methods, are scores assigned to each localized contig for the clustering, ordering and orienting stages of pseudomolecule construction. They provide a quantitative assessment of how well the clustering, ordering and orientation of a given contig is supported by the underlying Minimap2 alignments. These scores should be used, along with other homology analysis, as a quality control for RaGOO output. These scores can also be used to assess the efficacy of chimeric contig correction, or to check if an assembly diverges too much from the reference genome for accurate scaffolding. In general, if pseudomolecules pass these quality control checks, users can be more confident that RaGOO pseudomolecules are complete and representative of the underlying genome alignments.

## Conclusions

Our results show that RaGOO is a fast and accurate method for organizing genome assembly contigs into pseudomolecules. They also show that with a closely related reference genome, reference-guided scaffolding may yield substantially better scaffolding results than popular reference-free methods such as scaffolding with Hi-C data. In the process, we produced three tomato genome assemblies that are a valuable resource for the *Solanaceae* community and were selected to serve as the foundation for many additional tomato accessions we will be sequencing to establish a panSV-genome for use in biology and agriculture. For this purpose, the M82 assembly has already undergone extensive procedures to provide a complete and accurate assembly with an associated set of gene models and annotations.

## Methods

### Description of RaGOO Algorithm and Scoring Metrics

The complete RaGOO source code and documentation is available on github at https://github.com/malonge/RaGOO. RaGOO is written in Python3 and uses the python packages intervaltree and numpy, It also relies on Minimap2 that is available on github at https://github.com/lh3/minimap2. RaGOO also comes bundled with an implementation of Assemblytics for structural variation analysis.

#### Scaffolding algorithm overview

RaGOO utilizes alignments to a reference genome to cluster, order and orient contigs to form pseudomolecules. RaGOO internally invokes Minimap2, with k-mer size and window size both set to 19 bp, to obtain necessary mappings of contigs to a reference genome. By default, any alignments less than 1 kbp in length are removed. To cluster contigs, each contig is assigned to the reference chromosome which it covers the most. Coverage here is defined as the total number of reference chromosome base pairs covered in at least one alignment. Next, for each pseudomolecule group, the contigs in that group are ordered and oriented relative to each other. To do this, the longest (primary) alignment for each contig to its assigned reference chromosome is examined. Ordering is achieved by sorting these primary alignments by the start, then end alignment position in the reference. Finally, the orientation of that contig is assigned the orientation of its primary alignment. To produce pseudomolecules, ordered and oriented contigs are concatenated, with padding of “N” characters placed between contigs.

#### Scaffolding confidence scores

Each contig is assigned a confidence score for each of the 3 stages outlined above. The *clustering confidence score* is the number of base pairs a contig covered in its assigned reference chromosome divided by the total number of covered base pairs in the entire reference genome. To create a metric associated with contig ordering confidence, we defined a *location confidence*. First, the smallest and largest alignment positions, with respect to the reference, between a contig and its assigned reference chromosome are found. The location confidence is then calculated as the number of covered base pairs in this range divided by the total number of base pairs in the range. Finally, to calculate *orientation confidence*, each base pair in each alignment between a contig and its assigned reference chromosome casts a vote for the orientation of its alignment. The orientation confidence is the number of votes for the assigned orientation of the contig divided by the total number of votes.

#### Chimeric contig correction

Prior to clustering, ordering and orienting, RaGOO provides the option to break contigs which may be chimeric. RaGOO can identify and correct both interchromosomal and intrachromosomal chimeric contigs. Interchromosomal chimeric contigs are contigs which have significant alignments to two distinct reference chromosomes. To identify and break such contigs, all the alignments for a contig are considered. Alignments less than 10 kbp are removed, and the remaining alignments are unique anchor filtered [24]. If there are multiple instances where at least 5% of the total alignment lengths covered at least 100kbp of a distinct reference chromosome, a contig is deemed chimeric. To break the contig, alignments are sorted with respect to the contig start, then end positions, and the contig is broken where the sorted alignments transition between reference chromosomes.

Intrachromosomal chimeric contigs are contigs which have significant alignments to distance loci on the same reference chromosome. As with interchromosomal chimeric contigs, identification and breaking of intrachromosomal chimeric contigs starts with removing short and non-unique alignments. The remaining alignments are sorted with respect to the start, then end position in the reference chromosome. Next, the genomic distance between consecutive alignments is calculated, both with respect to the reference and the contig. If any of these distances exceeds user-defined thresholds, the contig is broken between the two alignments which the large distance between them.

### Simulated Reference-Guided Scaffolding

A simulated *S. lycopersicum* draft genome assembly was created by partitioning the Heinz SL3.0 reference genome, excluding chromosome 0, into scaffolds of variable length. Intervals along each chromosome were successively defined, with each interval length being randomly drawn from the distribution of observed M82 Canu contig lengths. Bedtools [42] was then used to retrieve the sequence associated with these intervals. Finally, simulated scaffolds with more than 50% “N” characters were removed and half of the remaining contigs were randomly reverse complemented. A second simulated assembly containing contigs, rather than scaffolds, was derived from these simulated scaffolds. Scaffolds were broken at any stretch of “N” characters longer than or equal to 20 bp, excluding the gap sequence. Any resulting contigs less than 10 kbp in length were also excluded. We call this pair of simulated assemblies the “easy” set of simulated data. To simulate a “hard” set of data, we started with the same “easy” scaffolds and added variation. To do this, we used SURVIVOR to simulate 10,000 indels ranging from 20 bp to 10 kbp in size. We also added SNPs at a rate of 1%. Again, we split these scaffolds into contigs resulting in a pair of “hard” simulated assemblies.

Given these “easy” and “hard” simulated scaffolds and contigs, RaGOO, Chromosomer, and MUMmer’s “show-tiling” utility were used for reference-guided scaffolding. For RaGOO, chimera breaking was turned off, and default parameters were used with the exception of the padding amount, which was set to zero. Chromosomer utilized Blast alignments with default parameters. Additionally, the “fragmentmap ratio” was set to 1.05 and the padding amount was set to zero. Show-tiling used default parameters. Since RaGOO and Chromosomer rely on aligners that allow for multithreading, both tools were run with 8 threads, while show-tiling was run with a single thread.

We recorded various measurements to evaluate the success of these tools in ordering and orienting simulated assemblies. Firstly, we observed the runtime, percentage of localized contigs and percentage of localized sequence. To assess clustering and orienting accuracy, we measure the percentage of localized contigs that had been assigned the correct cluster and orientation respectively. Finally, we used two measurements to assess ordering accuracy of each pseudomolecule. The first was the edit distance between the true and predicted order of contigs. This edit distance was normalized by dividing by the total number of contigs in the true ordering. The second ordering accuracy measurement was the percentage of correct adjacent contig pairs.

### Tomato Sequencing Data

#### Plant material and growth conditions

Seeds of the *S. lycopersicum* cultivar M82 (LA3475) were from our own stocks. Seeds of the *S. pimpinellifolium* accession BGV006775 were provided by E. van der Knaap, University of Georgia. Seeds of the *S. lycopersicum* breeding line Fla.8924 were from the stocks of S. Hutton, University of Florida. Seeds were directly sown and germinated in soil in 96-cell plastic flats and grown under long-day conditions (16-h light/8-h dark) for 21 days in a greenhouse under natural light supplemented with artificial light from high-pressure sodium bulbs (∼250 µmol m2 s1). Daytime and nighttime temperatures were 26– 28 **°**C and 18–20 **°**C, respectively, with a relative humidity of 40–60%.

#### Genome and transcriptome sequences

Genomic Illumina read data for BGV006775 were downloaded from the NCBI Sequence Read Archive (SRA) database (accession SRS3394566). Genomic Illumina read data for Fla.8924 (Lee et al., 2018) was provided by S. Hutton, University of Florida. Illumina read data for all transcriptomes were downloaded from ftp://ftp.solgenomics.net/user_requests/LippmanZ/public_releases/by_experiment/Park_etal/;[SeSo1] ftp://ftp.solgenomics.net/transcript_sequences/by_species/Solanum_lycopersicum/libraries/illumina/LippmanZ/[SeSo2] ; http://solgenomics.net/[SeSo3].[SeSo4][ZBL5]

#### Tissue collection and high molecular weight DNA extraction

For extraction of high molecular weight DNA, young leaves were collected from 21-day old light-grown seedlings. Prior to tissue collection, seedlings were incubated in complete darkness for 48h. Flash frozen plant tissue was ground using a mortar and pestle, and extracted in five volumes of ice-cold Extraction Buffer 1 (0.4 M sucrose, 10 mM Tris-HCl pH 8, 10 mM MgCl_2_, and 5 mM 2-Mercaptoethanol). Extracts were briefly vortexed, incubated on ice for 15 min, and filtered twice through a single layer of Miracloth (Millipore Sigma). Filtrates were centrifuged at 4,000 rpm for 20 min at 4 **°**C, and pellets were gently re-suspended in 1 ml of Extraction Buffer 2 (0.25 M sucrose, 10 mM Tris-HCl pH 8, 10 mM MgCl_2_, 1% Triton X-100, and 5 mM 2-Mercaptoetanol). Crude nuclear pellets were collected by centrifugation at 12,000 g for 10 min at 4 **°**C, and washed by re-suspension in 1 ml of Extraction Buffer 2 followed by centrifugation at 12,000 g for 10 min at 4 **°**C. Nuclear pellets were re-suspended in 500 μl of Extraction Buffer 3 (1.7 M sucrose, 10 mM Tris-HCl pH 8, 0.15% Triton X-100, 2 mM MgCl_2_, and 5 mM 2-Mercaptoethanol), layered over 500 μl Extraction Buffer 3, and centrifuged for 30 min at 16,000 g at 4 **°**C. Nuclei were re-suspended in 2.5 ml of Nuclei Lysis Buffer (0.2 M Tris pH 7.5, 2M NaCl, 50 mM EDTA, and 55 mM CTAB) and 1 ml of 5% Sarkosyl solution, and incubated at 60 **°**C for 30 min. To extract DNA, nuclear extracts were gently mixed with 8.5 ml of chloroform/isoamyl alcohol solution (24:1) and slowly rotated for 15 min. After centrifugation at 4,000 rpm for 20 min, ∼3 ml of aqueous phase was transferred to new tubes and mixed with 300 μl of 3M NaOAC and 6.6 ml of ice-cold ethanol. Precipitated DNA strands were transferred to new 1.5 ml tubes and washed twice with ice-cold 80% ethanol. Dried DNA strands were dissolved in 100 μl of Elution Buffer (10mM Tris-HCl, pH 8.5) overnight at 4 **°**C. Quality, quantity, and molecular size of DNA samples was assessed using Nanodrop (Thermofisher), Qbit (Thermofisher), and pulsed field gel electrophoresis (CHEF Mapper XA System, Biorad) according to the manufacturer’s’ instructions.

#### Nanopore library preparation and sequencing

DNA was sheared to 30kb using the Megarupter or 20kb using Covaris g-tubes. DNA repair and end-prep was performed using New England Biosciences kits NEBNext FFPE DNA Repair Kit and Ultra II End-Prep Kit. DNA was purified with a 1x AMPure XP bead cleanup. Next, DNA ligation was performed with NEBNext Quick T4 DNA Ligase, followed by another AMPure XP bead cleanup. DNA was resuspended in Elution buffer and sequenced according to the MinION standard protocol.

#### 10x Genomics library preparation and sequencing

1.12ng of high molecular weight gDNA was used as input to the 10X Genomics Chromium Genome kit v2 and libraries we prepared according to the manufacturer’s instructions. The final libraries, after shearing and adapter ligation, had an average fragment size of 626bp and were sequenced on an Illumina HiSeq, 2500 2×250bp.

#### Hi-C library preparation and sequencing

DNA extraction, library construction, and sequencing for Hi-C analyses was performed by Phase Genomics (Seattle, Washington) and conducted according to the supplier’s protocols. Young leaves from 21-day old light-grown and 48-h dark-incubated seedlings were wrapped in wet tissue paper and shipped on ice overnight.

### Initial *de novo* Assembly of Tomato Genomes

The Oxford Nanopore sequencing data for M82, BGV and FLA were assembled with Canu. For all three assemblies, default parameters were used with the expected genome size set to 950 Mbp. Assemblies were submitted to the UGE cluster at Cold Spring Harbor Laboratory for parallel computing. After assembly, it was determined that the M82 assembly contained bacterial contamination. To remove bacterial contigs from the assembly, the Canu contigs were aligned to all RefSeq bacterial genomes (downloaded on June 7, 2018) as well as the Heinz SL3.0 reference genome. If a contig covered more RefSeq bacterial genome base pairs than SL3.0 base pairs it was deemed a contaminant and removed from the assembly. In this paper “M82 Canu contigs” refers to the Canu contigs after contaminant contigs had been removed.

### Reference-Guided and Reference-Free Scaffolding of Tomato Genomes

The M82 Canu contigs were ordered and oriented into pseudomolecules with RaGOO, Chromosomer, and Nucmer’s ‘show-tiling’ utility. The Heinz SL3.0 reference, with chromosome 0 removed, was used for all tools. RaGOO used 8 threads with chimeric contig correction turned on and the gap padding size set to 200 bp. We also instructed RaGOO to skip 3 contigs which had low grouping accuracy scores. Chromosomer used 8 threads for BLAST alignments. The Chromosomer fragmentmap ratio was set to 1.05 and the gap padding size was set to 200 bp. Default parameters were used for show-tiling.

For reference-free scaffolding of the M82 assembly, 46,239,525,282 bp (∼60X coverage of the M82 Canu contigs) of 2×101 Hi-C sequencing reads were aligned to the M82 Canu contigs with BWA mem using the “-5” flag [43]. Aligned reads were then filtered with ‘samtools view’ to include alignments where both mates of a pair aligned as a primary, non-supplementary alignments (-F 2316) [44]. SALSA2 then utilized these alignments along with the M82 Canu assembly graph to build scaffolds. The SALSA2 “-m” flag was also set to “yes” in order to correct misassemblies in the M82 contigs and the expected genome size was set to 800 Mbp. Finally, we set “-e GATC” to correspond to the use of Sau3AI in the Hi-C library.

The structural accuracy of the M82 RaGOO pseudomolecules and SALSA2 scaffolds was assessed with dotplots and Hi-C density plots. For dotplots, both sequences were aligned to the Heinz SL3.0 reference (with chromosome 0 removed) with Minimap2 using the “-ax asm5” parameter. Alignments less than 12 kbp in length were excluded. For Hi-C visualization, the same Hi-C data described earlier was aligned to both sequences using the same parameters as were used for SALSA2. These alignments were then visualized with Juicebox [45]. Hi-C mates that mapped to the same restriction fragment were excluded from visualization.

Using the same parameters as M82, RaGOO was also used to order and orient the FLA and BGV Canu assemblies. BGV underwent two rounds of chimeric contig correction. Assemblytics structural variants for each assembly were compared with ‘SURVIVOR merge’, with the ‘max distance between breakpoints’ set to 1 kbp. Variants in chromosome 0 of the SL3.0 reference as well as variants which spanned more than 10% gaps were excluded from structural variant analysis.

### Tomato Genome Correction and Polishing

M82 RaGOO pseudomolecules were manually corrected for misassemblies and/or reference bias. Manual corrections were identified by visualizing Hi-C alignments to the M82 genome described in the previous sections. Firstly, three contigs with spurious alignments were removed from the pseudomolecules. Then, using Juicebox Assembly Tools, an inversion error was corrected on chromosome 3 and two ordering errors were corrected, one on chromosome 7 and one on chromosome 11. The “.assembly” file associated with these manual edits can be found in supplementary files. Gap filling and polishing was performed on the RaGOO pseudomolecules for the M82, FLA, and BGV tomato accessions. For each assembly, all respective Oxford Nanopore sequencing data used for assembly was used for gap filling with PBJelly.

After gap filling, we sought to find the most effective genome polishing strategy given our data. We used the gap-filled M82 assembly as a starting point for our tests. To polish this genome, we utilized the raw Oxford Nanopore data used for assembly as well as 10X Genomics Illumina Whole Genome Shotgun sequencing reads. We trimmed adapters and primers (23 bp from the beginning of read 1) and low-quality bases (40 bp from the ends of read 1 and read 2) from these 10X genomics data. With these data, we compared multiple polishing strategies using various alignment and polishing tools. First, we examined assemblies polished with or without Nanopolish [46]. For Nanopolish, the M82 raw Oxford Nanopore read set was aligned to the M82 assembly with Minimap2 using the “map-ont” parameter. Next, we compared assemblies polished with 1 or 2 rounds of Pilon polishing. For each round of polishing, the Illumina data was randomly subsampled to 40X coverage prior to alignment. Finally, we compared bwa mem, Bowtie2 and ngm for short-read alignment prior to Pilon polishing [47, 48]. We used bwa mem and ngm with default parameters, while Bowtie2 was run with the “--local” parameter.

We used MUMmer’s ‘dnadiff’ utility to compare the efficacy of these polishing pipelines (**Supplementary Table 4**). For dnadiff analysis, polished assemblies and the SL3.0 reference were broken into contigs by breaking sequences at gaps of 20 bp or longer. Then, assemblies were aligned to the reference contigs with nucmer using the ‘-l 100 -c 500 -maxmatch’ parameters. After determining that 2 rounds of Pilon polishing with Bowtie2 yielded the best results, we applied the same pipeline to the BGV and FLA assemblies using ∼23 × coverage and ∼26X coverage of paired-end Illumina short-read data was used for BGV and FLA respectively. BUSCO was used to evaluate genome completeness of the polished M82, BGV, and FLA assemblies. The Solanaceae odb10 database was used with the “species” parameter set to “tomato”.

### Tomato Genome Annotation

We annotated protein coding genes in the M82, FLA and BGV assembly using the Maker v3.0 pipeline on Jetstream by providing repeats, full length cDNA sequences and proteins from Heinz 1706 ITAG3.2 assembly [49]. Simple, low complexity and unclassified repeats were excluded from masking. We additionally provided Maker with an M82 reference transcriptome derived from 50 M82 RNA-seq libraries. RNA-Seq reads were aligned to the M82 genome using STAR, a splice aware aligner [50]. These alignments were used to assemble transcripts and establish a consensus transcriptome using StringTie and TACO Respectively [51, 52]. We ran Maker using parameters est2genome set to 1, protein2genome set to 1 and keep_preds set to 1 to perform the gene annotation. Low consensus gene models with an AED score above 0.5 were filtered from the Maker predicted gene models. We additionally removed gene models shorter than 62 bp folllowing the cutoffs used for the ITAG3.2 annotation. Putative gene functions were assigned to the MAKER gene models via Interproscan protein signatures and blastp protein homology search [53]. blastp queried the UniProtKB/Swiss-Prot and Heinz 1706 ITAG3.2 protein databases, filtering out alignments with an e-value greater than 1e-05 [54]. We further filtered out genes that did not have an associated gene function in either Interproscan, UniprotKB/Swiss-Prot or ITAG3.2.

### Arabidopsis Structural Variant Analysis

The 1001 genomes database was mined for accessions for which there was at least 50X coverage of paired end sequencing data. We also required that the read length be at least 100 bp. For practical reasons, we excluded accessions with excessive coverage. For each of the remaining accessions, the fastq files were randomly subsampled in order to achieve exactly 50X coverage. Subsampled reads were then assembled with the SPAdes assembler, with k-mer size set to 33, 55, 77, and 99, and otherwise default parameters. These draft assemblies were then ordered and oriented with RaGOO using default parameters and the TAIR 10 reference genome (GCA_000001735.1). RaGOO also provided structural variants, with the minimum variant size set to 20bp. Of the chromosome-scale assemblies, a few assemblies with a genome size greater than 150 Mbp were removed due to putative sample contamination. After this filtering, assemblies and structural variant calls for 103 accessions remained.

Variants that were called in chromosome 0 or the chloroplast/mitochondrial chromosomes were discarded. Also, variants which had more than a 10% overlap with a gap were excluded. To find unique variants across multiple samples, SURVIVOR merge was used such that a variant only had to be present in at least one sample for it to be reported. Therefore, given all 103 samples, this yielded the union of all variants present in the pan-genome. To find shared variants across multiple samples, SURVIVOR merge was used such that a variant must have been present in all samples to be reported. This effectively provided the intersection of variants in the pan-genome. In all instances of using SURVIVOR merge, the ‘max distance between breakpoints’ was set to 1 kbp. Also, the strand of the SV was taken into account, while distance based on the size of the variant was not estimated. Finally, the minimum variant size was set to 20 bp to be consistent with the RaGOO parameters. Bedtools was used to find variant/gene intersections.

## Data Availability

Sequencing data, genome assemblies, annotations and structural variation calls for all samples are available at http://share.schatz-lab.org/ragoo/. The tomato genomes are also available in the Sol Genomics Network (https://solgenomics.net/)

## Author Contributions

MA designed and wrote the RaGOO software. MA and MS conceived of and executed experiments with simulated data. SS, SG, XW, and ZL chose tomato accessions for genome assembly, collected plant tissue, and collected sequencing data. MA and MS assembled draft tomato genomes and removed contaminant contigs. MA and MS conceived of and executed experiments to compare reference-free and reference-guided scaffolding of the M82 genome. MA finished the tomato genomes via manual correction with Hi-C, gap filling, and polishing. SR and MS performed gene finding an annotation in the M82 genome and assessed the quality of the results. MA, FS, and MS conceived of an executed *A. thaliana* pan-genome analysis. MA and MS wrote the majority of the manuscript, with all other authors making contributions. All authors read and approved the final manuscript.

## Acknowledgements

We thank Esther van der Knaap and Sam Hutton for providing seed stocks and thank W. Richard McCombie for helpful discussions. This work was supported by the National Science Foundation (DBI-1350041 and IOS-1445025 to M.C.S.; IOS-1732253 to M.C..S and Z.B.L.) and the US National Institutes of Health (R01-HG006677 to M.C.S. and UM1 HG008898 to F.J.S.). We also thank the NSF XSEDE project for providing compute resources on JetStream for annotation (project MCB180087 to M.C.S.)

## Figures

**Supplementary Figure 1.**
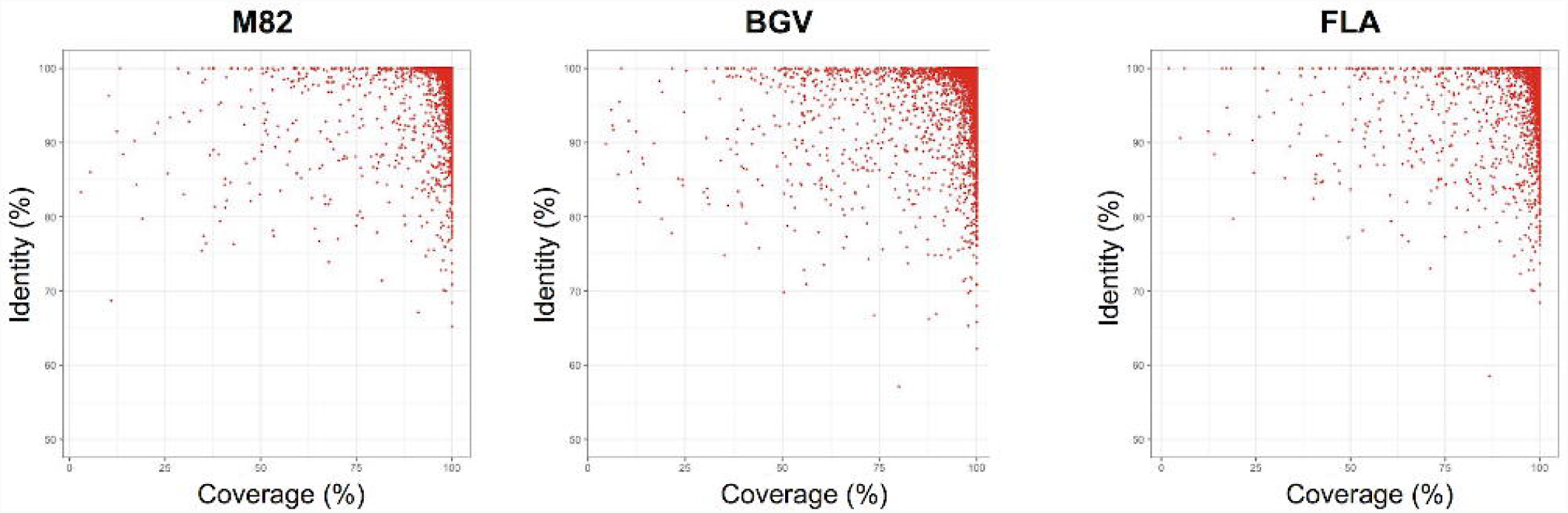
Heinz cDNA Alignment. Coverage vs. identity of GMAP alignments of ITAG3.2 cDNA to the M82, BGV, and FLA assemblies.

**Supplementary Figure 2.**
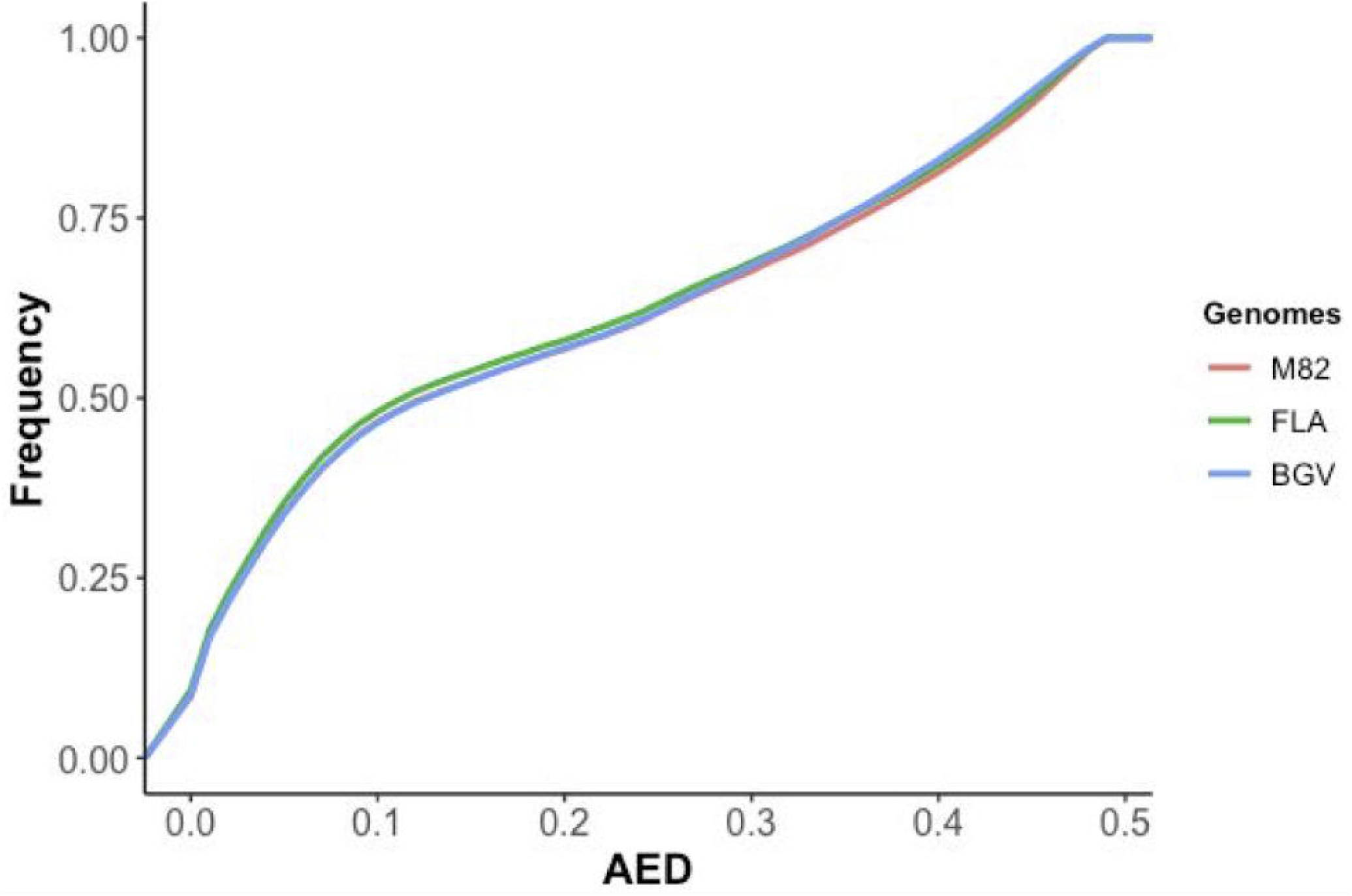
Annotation Edit Distances. Cumulative Distribution of Annotation Edit Distance (AED) in Maker annotated Genomes.

**Supplementary Figure 3.**
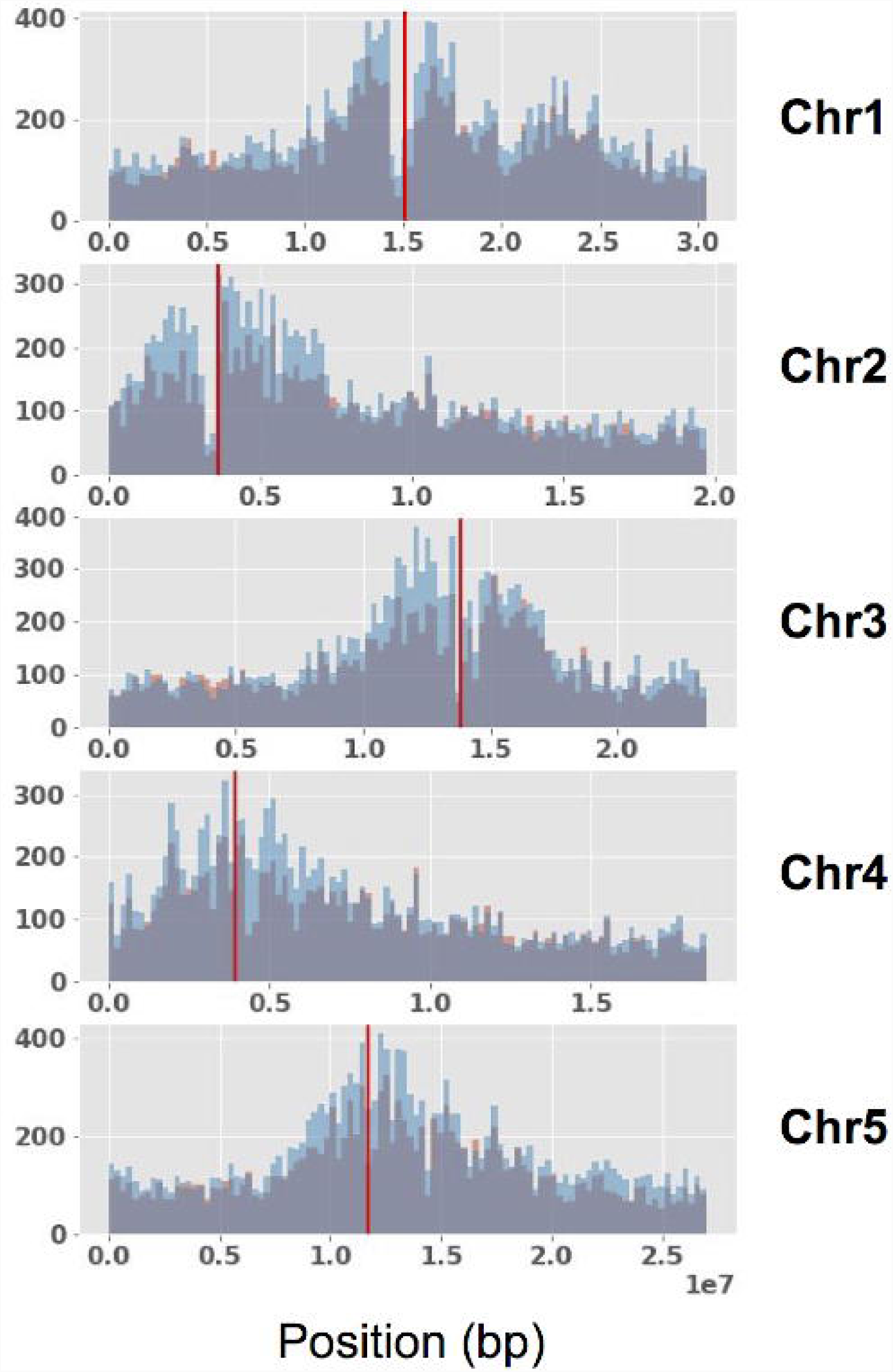
A. thaliana PanSV-Genome SV Distribution. Distribution of insertions (orange) and deletions (blue) along each of the 5 Arabidopsis thaliana chromosomes. The red vertical lines indicate the centromere midpoints.

